# Activation of SnRK2 by the Raf-like kinase ARK represents a common mechanism of ABA response in embryophytes

**DOI:** 10.1101/2020.04.03.024448

**Authors:** Mousona Islam, Takumi Inoue, Mayuka Hiraide, Nobiza Khatun, Akida Jahan, Keiko Kuwata, Taishi Umezawa, Izumi Yotsui, Yoichi Sakata, Daisuke Takezawa

**Affiliations:** Graduate School of Science and Engineering, Saitama University, Shimo-ohkubo 255, Sakura-ku, Saitama 338-8570, Japan; Plant Tissue Culture Section, Biological Research Division, Bangladesh Council of Scientific and Industrial Research (BCSIR), Dhaka 1205, Bangladesh; Institute of Transformative Bio-Molecules (WPI-ITbM), Nagoya University, Furo-cho, Chikusa-ku, Nagoya, 464-8601, Japan; Graduate School of Bio-Applications and Systems Engineering, Tokyo University of Agriculture and Technology, 2-24-16 Nakacho, Koganei City, Tokyo 184-8588, Japan; Department of Bioscience, Tokyo University of Agriculture, 1-1-1 Sakuragaoka, Setagaya-ku, Tokyo 156-8502, Japan

**Keywords:** abiotic stress, abscisic acid, cold, *Physcomitrella patens*, protein kinase

## Abstract

The Raf-like protein kinase ARK previously identified in the moss *Physcomitrella patens* acts as an upstream regulator of subgroup III SnRK2, the key regulator of abscisic acid (ABA) and abiotic stress responses. However, the mechanisms underlying activation of ARK by ABA and abiotic stress for the regulation of SnRK2 including the role of ABA receptor-associated group A PP2C (PP2C-A) are not understood. We identified Ser1029 as the phosphorylation site in the activation loop of ARK, which provided a possible mechanism for regulation of its activity. Analysis of transgenic *ark* lines expressing ARK-GFP with Ser1029-to-Ala mutation indicated that this replacement causes reductions in ABA-induced gene expression, stress tolerance and SnRK2 activity. Immunoblot analysis using an anti-phosphopeptide antibody indicated that ABA treatments rapidly stimulate Ser1029 phosphorylation in wild type. The phosphorylation profile of Ser1029 in ABA-hypersensitive *ppabi1* lacking PP2C-A was similar to that in wild type, whereas little Ser1029 phosphorylation was observed in ABA-insensitive *ark* missense lines. Furthermore, newly isolated *ppabi1 ark* lines showed ABA-insensitive phenotypes similar to those of *ark* lines. These results indicate that ARK is a primary activator of SnRK2, preceding negative regulation by PP2C-A in bryophytes, which provides a prototypal mechanism for ABA and abiotic stress-responses in embryophytes.

**One sentence summary:** Physiological characterization of various moss mutants revealed a common mechanism for phytohormone responses under water deficit in all land plants.

## INTRODUCTION

The phytohormone abscisic acid (ABA) regulates a variety of developmental as well as physiological processes in plants. In vegetative tissues, elevated endogenous ABA in response to water deficit in shoots contributes to stomatal closure and tolerance to environmental abiotic stresses such as drought and cold (Rock *et al*., 2010). Studies on Arabidopsis have revealed that ABA elicits these cellular responses by binding to the intracellular ABA receptor PYRABACTIN RESISTANCE1 (PYR1)/PYR1-LIKE/REGULATORY COMPONENTS OF ABA RECEPTORS (PYR/PYL/RCAR) (Park *et al*., 2009; Ma *et al*., 2009). The ABA-PYR/PYL/RCAR complex binds to group A protein phosphatase 2C (PP2C-A) to inhibit its phosphatase activity, which leads to activation of subclass III SNF1-related protein kinase2 (SnRK2) (Umezawa *et al.,* 2009; Vlad *et al.,* 2009; Nishimura *et al*., 2010; Rodrigues *et al*., 2013). Of the three SnRK2 subgroups, subclass III SnRK2 is recognized as the key regulator of ABA signaling since it phosphorylates various cellular substrates including bZIP transcription factors that activate a number of ABA-induced genes (Marcotte *et al*., 1989; Uno *et al*., 2000; Fujita *et al*., 2005). Activity of subclass III SnRK2 is stimulated by ABA-induced phosphorylation at specific Ser residues in the “activation loop” located between subdomains VII and VIII of the kinase domain (Boudsocq *et al*., 2004; Konrev *et al*., 2006; Boudsocq *et al*., 2007). This phosphorylation event is operated by either autophosphorylation by SnRK2 itself or by other protein kinases, though detailed mechanisms have not been clarified.

The roles of ABA in stress responses have been demonstrated not only in angiosperms but also in basal embryophytes, mosses and liverworts. Protonemata of the moss *Physcomitrella patens* acquire tolerance to desiccation, hyperosmosis and freezing upon exogenous ABA treatment (Minami *et al*., 2003; Khandelwal *et al*., 2010; Koster *et al*., 2010). ABA induces expression of genes for late embryogenesis abundant (LEA)-like proteins (Cuming and Lane, 1979) with boiling-soluble characteristics mediated by conserved ABA-responsive promoter elements in the moss protonemata (Knight *et al*. 1995). The *P. patens* genome has four genes for putative PYR/PYL/RCAR, two genes for PP2C-A and four genes for subclass III SnRK2 but no SnRK2 of other subclasses (Rensing *et al*., 2008; Sakata *et al*., 2014). Disruption of genes for PP2C-A (*PpABI1A* and *PpABI1B*) in the *ppabi1* line resulted in an ABA-hypersensitive response and constitutive desiccation tolerance (Komatsu *et al*., 2013). Furthermore, disruption of all four subgroup III SnRK2 genes (*PpSnRK2A* to *PpSnRK2D*) resulted in a loss of ABA sensitivity (Shinozawa *et al*., 2019). These findings suggest that core signaling mechanisms for ABA signaling are common in embryophytes including bryophytes and angiosperms.

While the role of PP2C-A in negative regulation of subgroup III SnRK2 has been well documented, the molecules responsible for positive regulation of SnRK2 have not been clarified. By analysis of a *P. patens* mutant designated AR7 that shows reduced SnRK2 activity, we previously reported that ABA- and abiotic stress-responsive Raf-like kinase (ARK) plays a crucial role in integration of ABA and abiotic stress response (Minami *et al*., 2006; Saruhashi *et al*., 2015). ARK with 1148 amino acids consists of the C-terminal protein kinase domain with similarity to group B3 Raf-like protein kinase (B3-Raf) and a large non-kinase region with unknown function toward the N-terminus. AR7 was not only insensitive to ABA but also less responsive to hyperosmosis and cold, indicating that ARK might play a role in the integration of ABA and these abiotic signals. AR7 has a missense mutation in Ser532 changed to Phe in the non-kinase region of ARK, suggesting that the region toward the N-terminus to the kinase domain might play a role in the regulation of ARK activity. Null mutations of *ARK* causing loss of ABA-induced desiccation tolerance have also been reported (Yasumura *et al*., 2015; Stevenson *et al*., 2016). The kinase domain of ARK fused to glutathione-S-transferase phosphorylated and activated subclass III SnRK2 (PpSnRK2B and PpSnRK2D) of *P. patens in vitro*, indicating that ARK acts as a positive regulator of SnRK2 in the ABA signaling process in bryophytes (Saruhashi *et al*., 2015; Shinozawa *et al*., 2019).

Although our studies have indicated that ARK is one of key regulators of abiotic stress signaling, little is known about how ARK activity is regulated by ABA. Studies on various eukaryotic Raf-related protein kinases highlighted phosphorylation-mediated activation of the kinases and the role of their N-terminal domains in the regulation of phosphorylation (Köhler and Brummera, 2016). By phosphopeptide mapping of *P. patens* ARK, we showed that ARK is phosphorylated in the activation loop of the kinase domain, providing a possible mechanism for regulation of ARK (Saruhashi *et al*., 2015). In the present study, we therefore focused on phosphorylation and activation of ARK during ABA and stress response using *P. patens* lines for which site-specific mutations have been introduced. The results of our analysis indicated that ARK phosphorylation in the activation loop is critical for SnRK2 regulation during ABA response. We also analyzed changes in ARK phosphorylation during ABA and abiotic stress treatments using an anti-phosphopeptide antibody in various mutants as well as in WT *P. patens*.

## RESULTS

### Analysis of ARK-GFP lines with mutations in putative phosphorylation sites

Our previous study showed by phosphopeptide mapping that Ser1029 and Ser1030 in the activation loop of the ARK kinase domain are putative phosphorylation sites (Saruhashi *et al*., 2015). To determine which serine residues are critical for activation of ARK, ARK-GFP constructs with or without mutations in these phosphorylation sites (ARK^S1029A^-GFP ARK^S1030A^-GFP ARK^S1029A/S1030A^-GFP) (Fig. 1A) were generated and introduced into AR7 with S532F mutation in ARK. Growth tests on a medium containing 10 μM ABA indicated that protonemal growth of AR7 and AR7 expressing non-mutated ARK-GFP and ARK^S1030A^-GFP was severely inhibited by ABA. In contrast, growth of AR7 expressing ARK^S1029A^-GFP and ARK^S1029A/S1030A^-GFP was less sensitive to ABA. We also generated AR7 expressing ARK^S532F^-GFP for comparison. As expected, there was little inhibition of growth of AR7 expressing ARK^S532F^-GFP by ABA (Fig. 1B, Fig. S1).

**Figure 1:**
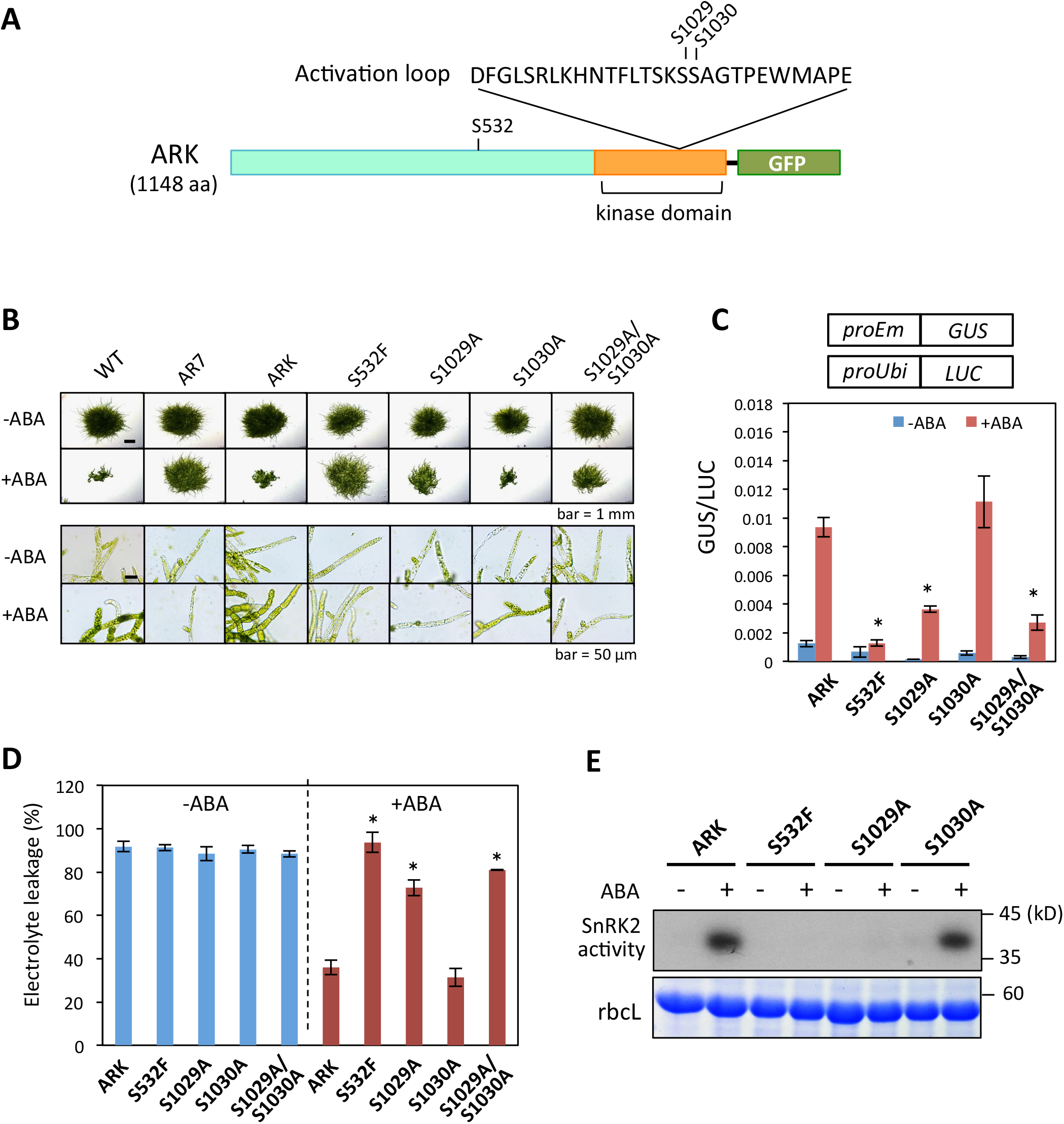
Analysis of *Physcomitrella patens* lines expressing ARK-GFP constructs. (**A**) Schematic representation of ARK-GFP showing positions of Ser532 (S532) in the non-kinase domain and putative phosphorylation sites, Ser1029 (S1029) and Ser1030 (S1030), located in the activation loop of the kinase domain. (**B**) Growth responses to ABA of wild type (WT), AR7, and AR7 lines expressing ARK-GFP without mutation (ARK) and with S532F, S1029A, S1030A and S1029A/S1030A mutations. Protonemata of these lines were cultured with or without 10 μM ABA for one week. ABA-induced gene expression in transgenic lines. Protonemata were bombarded with plasmid constructs of the *Em* promoter fused with the *beta-glucuronidase* (*proEm-GUS*) gene and the rice *ubiquitin* promoter fused with the *luciferase* gene (*proUbi-LUC*) and were cultured with or without 10 μM ABA for one day before GUS and LUC assays. Error bars indicate standard error of the mean. \**p* < 0.05 in the t-test (n = 3) compared with the ABA-treated ARK line. (**D**) Freezing tolerance of ARK-GFP lines. Protonemata were cultured with or without 10 μM ABA for one day and subjected to freezing at 10°C. After thawing, electrolyte leakage was determined to estimate the damage caused by freezing. Error bars indicate standard error of the mean. \**p* < 0.05 in the t-test (n = 3) compared with the ABA-treated ARK line. (**E**) In-gel kinase assays for detection of SnRK2 activity. Proteins extracted from protonemata were electrophoresed using SDS-polyacrylamide gel polymerized with histone IIIS as a substrate. After denaturation and renaturation processes, the proteins were reacted with ^32^P-ATP and radioactive signals were detected by exposure to X-ray film. Staining of the large subunit of ribulose bisphosphate calboxylase (rbcL) is shown as a control.

Using these lines, we analyzed ABA-induced gene expression by transient assays using the beta-glucuronidase (*GUS*) reporter gene fused to the ABA-inducible *Em* promoter (*proEm*) (Knight *et al*., 1995; Sakata *et al*., 2010). The results of assays indicated that AR7 lines expressing ARK^S1029A^-GFP and ARK^S1029A/S1030A^-GFP showed only a partial ABA response compared with the lines with ARK-GFP and ARK^S1030A^-GFP (Fig. 1C). We also analyzed ABA-induced freezing tolerance in these lines. We previously showed that ABA treatment of protonemata increases freezing tolerance of the cells in WT but not in AR7 (Minami *et al*., 2006; Bhyan *et al*., 2011). Tests for freezing tolerance indicated that AR7 lines with ARK-GFP and ARK^S1030A^-GFP showed similar levels of freezing tolerance induced by ABA treatment, whereas the lines with ARK^S1029A^-GFP and ARK^S1029A/S1030A^-GFP showed only a slight enhancement in the tolerance induced by ABA treatment (Fig. 1D). These results indicated that the mutation in Ser1029 to Ala causes reductions in ABA response for gene expression and stress tolerance.

Since ARK is postulated to be an activator of SnRK2, we carried out in-gel kinase assays of ARK-GFP mutant lines using a gel containing histone IIIS, by which we previously showed that AR7 has little SnRK2 kinase activity (Saruhashi *et al*., 2015). The results of assays indicated that the lines with ARK^S532F^-GFP and ARK^S1029A^-GFP showed little SnRK2 activity, while the ARK^S1030A^-GFP line showed ABA-induced SnRK2 activity similar to that of the ARK-GFP line (Fig. 1E). These results indicated that Ser1029 of ARK might be a critical phosphorylation site for activation of SnRK2 in *P. patens*.

To determine the importance of Ser1029 phosphorylation, anti-phosphopeptide antibody specific to Ser1029 phosphorylation (anti-P-S1029) was raised and used for immunoblot analysis (Fig. 2A). We detected immuno-reacting signals around the expected molecular mass of ARK-GFP (157 kDa) that were enhanced by 15-min treatment with 10 μM ABA in the non-mutated ARK-GFP line. Similar enhancement of signals by ABA was also detected in the ARK^S1030A^-GFP lines. In contrast, the signals were very faint in ARK^S532F^-GFP and ARK^S1029A^-GFP lines (Fig. 2A). Phosphopeptide analysis of ARK-GFP proteins immunoprecipitated from ABA-treated protonemata using anti-GFP antibody microbeads indicated that Ser phosphorylation in the peptide of amino acids 1029-1043 occurred in the ARK-GFP and ARK^S1030A^-GFP lines but not in the ARK^S1029A^-GFP and ARK^S532F^-GFP lines (Table S2). The anti-P-S1029 antibody was also used for detection of phosphorylation of native ARK (Fig. 2B). The signal detected at around 130 kilodaltons by the anti-P-S1029 antibody was increased after 15 min of ABA treatment, while the levels of ARK polypeptide detected by an antibody against ARK C-terminal peptide (anti-ARKc) were unchanged. The level of Ser1029 phosphorylation was highest at 30 min and gradually decreased with time, but the enhanced phosphorylation was still observed even after 3 h (Fig. S2). We also found that osmotic treatment with 0.5 M mannitol stimulates the Ser1029 phosphorylation, though to a lesser extent than that with ABA treatment (Fig. S3).

**Figure 2:**
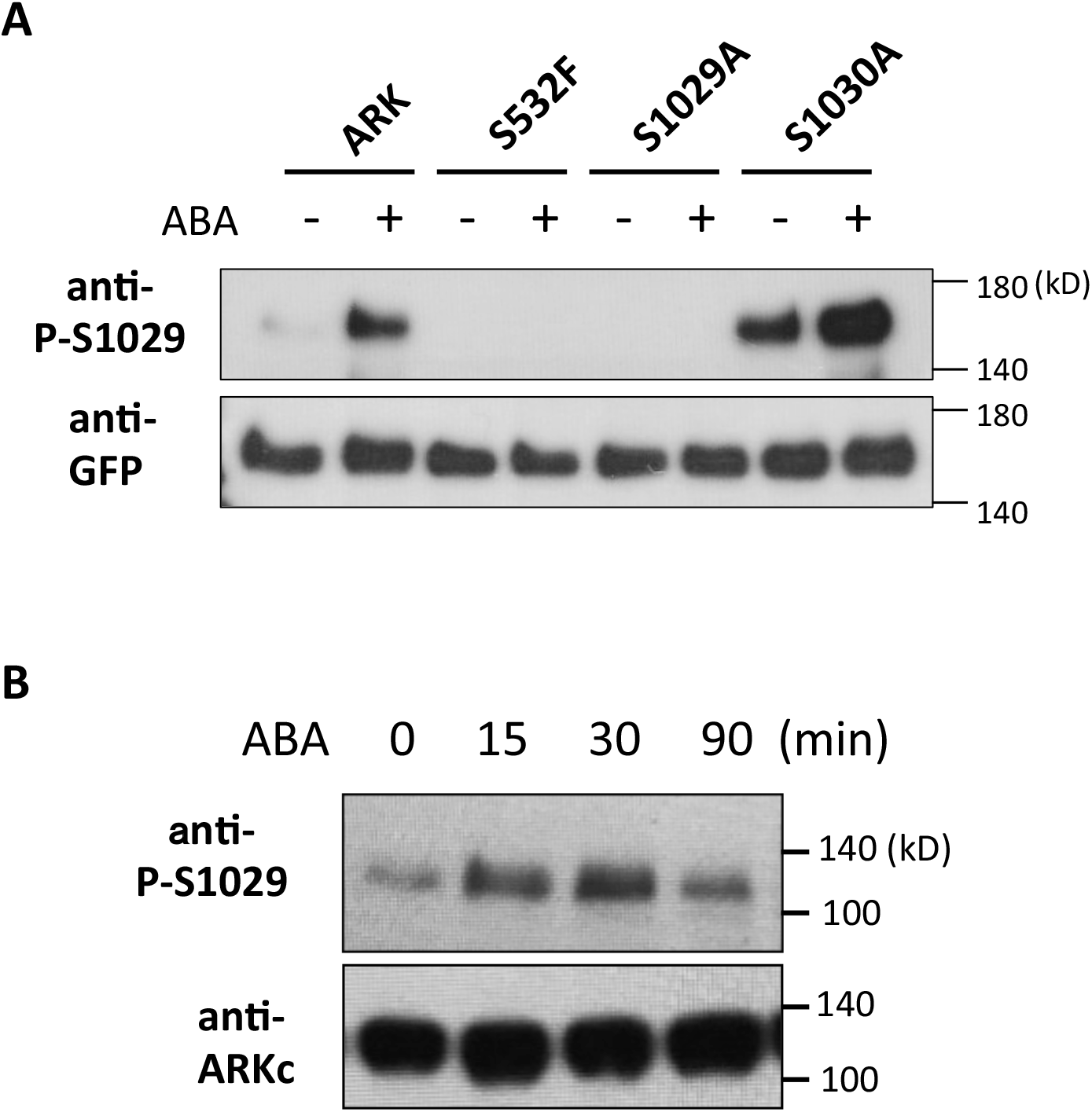
ABA-stimulated phosphorylation of Ser1029 detected by anti-phosphopeptide antibody. (**A**) Proteins from protonemata of transgenic *Physcomitrella patens* AR7 lines expressing ARK-GFP (ARK) and ARK-GFP with S532F, S1029A, and S1030A mutations with or without treatment with 10 μM ABA for 15 min were subjected to immunoblot analysis using an anti-phospho-Ser1029 (P-S1029) antibody and an anti-GFP antibody. Since levels of ARK-GFP accumulation vary among these lines, the amount of proteins loaded per lane was adjusted using the anti-GFP antibody. Note that ARK^S1030A^-GFP tends to give enhanced signals for an unknown reason. (**B**) Detection of native ARK in WT *P. patens*. Protonemata were treated with 10 μM ABA for the indicated time periods, and extracted proteins were reacted with the anti-P-S1029 antibody and an antibody that recognizes the C-terminal 15-amino-acid peptide of ARK (anti-ARKc).

### Analysis of AR144 (*ark^D995N^*) with mutation in the kinase catalytic core

The fact that AR7 with the S523F mutation in the non-kinase region but with an intact kinase domain showed little Ser1029 phosphorylation and ABA response raised the question about whether Ser1029 phosphorylation requires catalytic activity of ARK itself. To determine the role of the kinase domain of ARK, we searched for new *ark* alleles by screening of more than 6 x 10^6^ mutagenized protonema cells, and we isolated the ABA-insensitive AR144 line, in which Asp995 in the kinase domain of ARK has been changed to Asn (Fig. 3A). This Asp residue corresponds to a highly conserved Asp in the “HRD motif” in the catalytic loop of various eukaryotic protein kinases (Fig. 3A). This Asp residue is essential for the kinase activity since it is assigned to be responsible for correct orientation of the target hydroxyl group of Ser in the substrate peptide (Kannan and Neuwald 2005; Kornev *et al*., 2006).

**Figure 3:**
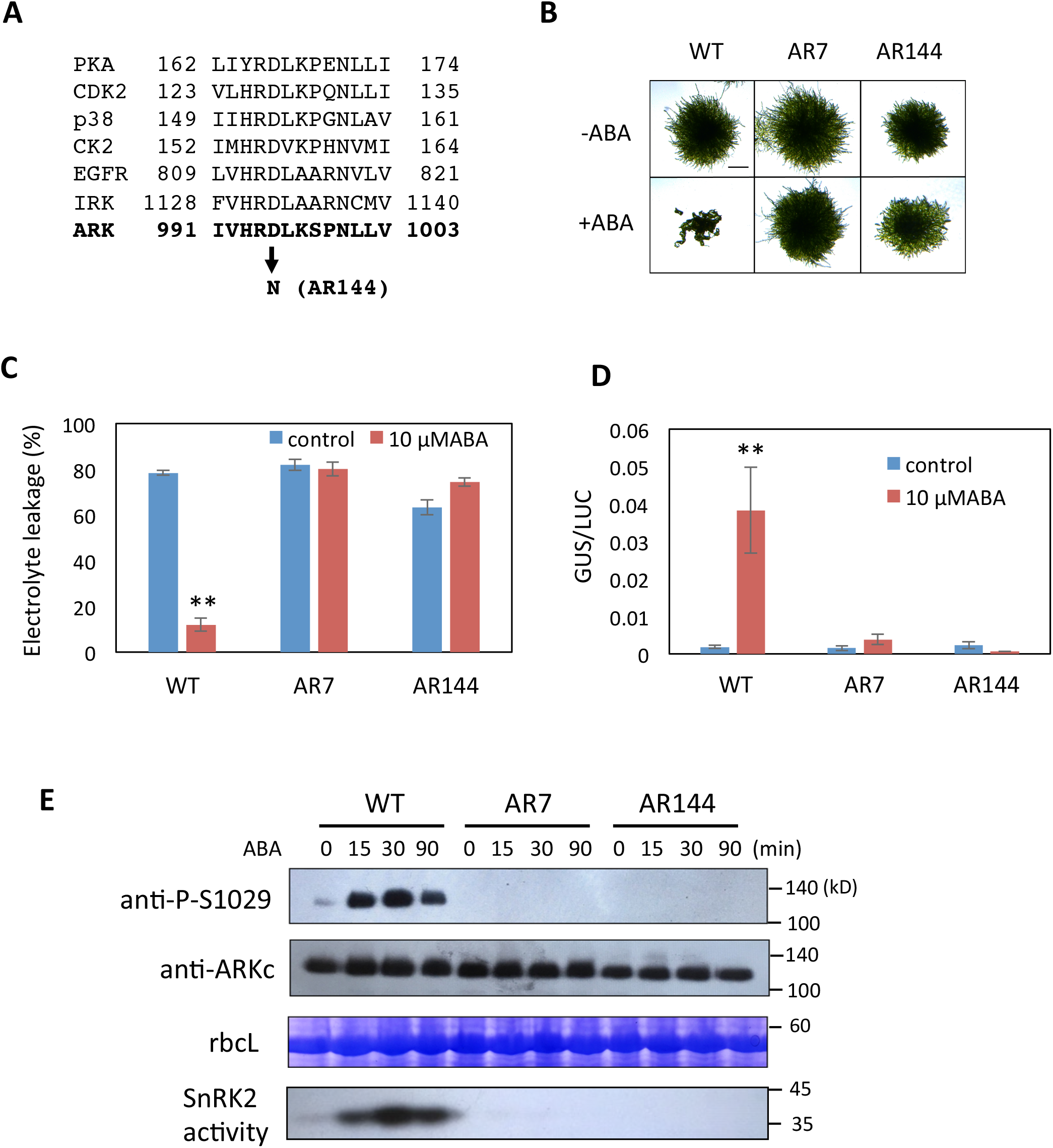
ABA response of AR144 of *Physcomitrella patens* with a mutation in the catalytic core of the kinase domain of ARK. (**A**) Comparison of amino acid sequences in the catalytic core of protein kinase A (PKA), cyclin-dependent protein kinase2 (CDK2), p38, casein kinase2 (CK2), and epidermal growth factor receptor (EGFR) (adapted from Nolen *et al*., 2004) with ARK. The conserved Asp (D) has been changed to Asn (N) in AR144. (**B**) Growth responses to ABA. Protonemata of WT, AR7 and AR144 were grown on a medium with 10 μM ABA for one week. (**C**) Comparison of freezing tolerance. Protonemata incubated with or without 10 μM ABA for one day were frozen to −4°C. After hawing, electrolyte leakage was measured to estimate the extent of freezing injury. Error bars indicate standard error of the mean. Transient gene expression assays of WT, AR7 and AR144. Protonemata were bombarded with plasmid constructs of *proEm-GUS* and *proUbi-LUC* and were cultured with or without 10 μM ABA for one day for GUS and LUC assays. Error bars indicate standard error of the mean. \*\**p* < 0.01 in the t-test (n = 3) compared with control WT without ABA treatment. (**E**) Effect of ABA on Ser1029 phosphorylation in WT, AR7 and AR144. Protonemata were treated with 10 μM ABA for different times and used for immunoblot analysis with anti-phospho-Ser1029 (P-S1029) and anti-ARKc antibodies. Staining of the large subunit of ribulose bisphosphate calboxylase (rbcL) is shown as a control. Results of an in-gel kinase assay for detection of SnRK2 are also shown.

When grown on a medium containing 10 μM ABA, protonemata of AR144 showed ABA-insensitive growth similar to that of AR7 (Fig. 3B). In freezing tolerance tests, freezing-induced leakage of electrolytes in WT protonemata was remarkably reduced by pretreatment with ABA, whereas the leakage was not mitigated by ABA in AR144 protonemata as in the case of AR7 (Fig. 3C). Transient reporter assays indicated that there was no ABA-increased gene expression in AR144 (Fig. 3D) in contrast to AR7 with an intact kinase domain showing a slight increase. We analyzed phosphorylation of ARK in AR144 by using the anti-P-Ser1029 antibody to examine whether the ARK kinase activity is necessary for its phosphorylation and is correlated with activation of SnRK2. The results of analyses indicated that while both Ser1029 phosphorylation and SnRK2 activity were enhanced by ABA in WT, neither Ser1029 phosphorylation nor SnRK2 activity was detected in AR144 as well as in AR7 (Fig. 3E). These results indicated that the catalytic activity of ARK is necessary for Ser1029 phosphorylation and cellular ABA response.

### Effect of cold on ARK phosphorylation

We previously reported that the freezing tolerance of *P. patens* protonema cells is enhanced by exposure to low temperatures (Minami *et al*., 2005). This cold acclimation process accompanied accumulation of LEA-like proteins that mitigate freezing damage (Minami *et al*., 2005; Sasaki *et al*., 2014). Comparison of freezing-induced electrolyte leakage in WT and AR7 indicated that cold treatment (0°C) enhanced freezing tolerance in WT but only slightly in AR7 (Fig. 4A), correlated with accumulation of the LEA-like 17B9 protein in boiling-soluble fractions (Fig. 4B). Immunoblot analysis of WT and AR7 protonemata with anti-P-S1029 and anti-ARKc antibodies revealed that phosphorylation of Ser1029 was enhanced in WT with increased abundance of ARK polypeptide during cold acclimation (Fig. 4B). The Ser1029 phosphorylation was correlated with enhancement of SnRK2 activity. In AR7, however, little enhancement of both Ser1029 phosphorylation and SnRK2 activity was observed during cold acclimation, although cold-enhanced accumulation of ARK polypeptide was observed. These results suggest that Ser1029 phosphorylation of ARK plays a role in the cold acclimation process in the moss cells.

**Figure 4:**
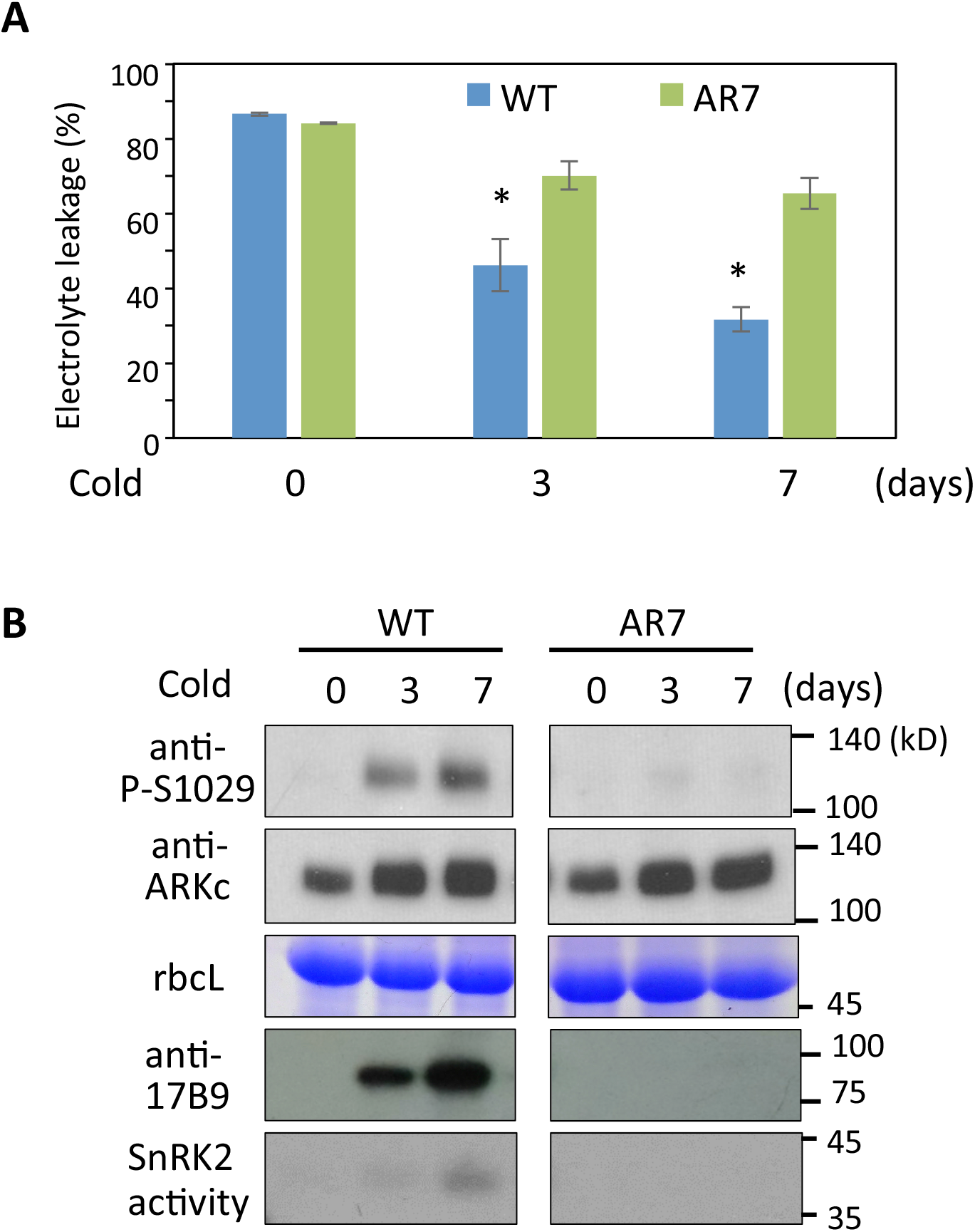
Cold responses mediated by ARK in *Physcomitrella patens*. (**A**) Changes in freezing tolerance during cold acclimation in wild type (WT) and AR7 (*ark^S532F^*). Protonemata that had been cold-acclimated for the indicated days were frozen to −3°C and injury rate was estimated by measurement of electrolyte leakage after thawing. Error bars indicate standard error of the mean. \**p* < 0.05 in the t-test (n = 3) compared with non-acclimated protonemata of each line. (**B**) Immunoblot analysis with anti-phospho-Ser1029 (P-S1029) and anti-ARKc antibodies. Coomassie Brilliant Blue staining of the large subunit of ribulose bisphosphate calboxylase (rbcL), results of immunoblot analysis for cold-induced 17B9 protein, and SnRK2 activity analyzed by an in-gel kinase assay are also shown.

### Roles of PP2C-A in regulation of ARK phosphorylation and SnRK2 activity

PP2C-A associated with the PYR/PYL/RCAR receptor is thought to be a critical regulator of SnRK2 in Arabidopsis (Park *et al*., 2009; Umezawa *et al*., 2009). Komatsu *et al*. (2013) reported that the *P. patens ppabi1* line lacking both of the two genes for PP2C-A showed desiccation tolerance without ABA treatment. Protonemata of *ppabi1* frequently form spherical “brood cells” (Arif *et al*., 2019) under normal growth conditions, while these cells are formed by ABA treatment for several days in WT protonemata (Fig. 5A). Interestingly, the *ppabi1* protonemata are still responsive to ABA and can form more brood cells upon ABA treatment (Fig. 5A). In-gel kinase assays of *ppabi1* indicated that ABA-dependent SnRK2 activity in *ppabi1* was greater than that in WT, though the activity remained low without exogenous ABA (Komatsu *et al*., 2013), indicating that PP2C-A is not the only regulator of SnRK2. Since ARK has been identified as a positive regulator of SnRK2, we suspected that PP2C-A might affect SnRK2 through regulation of ARK. Comparison of ABA-induced changes in Ser1029 phosphorylation of ARK and SnRK2 activity in the same plant extract indicated that phosphorylation of ARK was stimulated by ABA in *ppabi1* in a manner similar to that of WT, in contrast to a remarkable enhancement of SnRK2 activity in *ppabi1* (Fig. 5B). These results indicated that PP2C-A is not required for ABA-induced activation of ARK phosphorylation.

**Figure 5:**
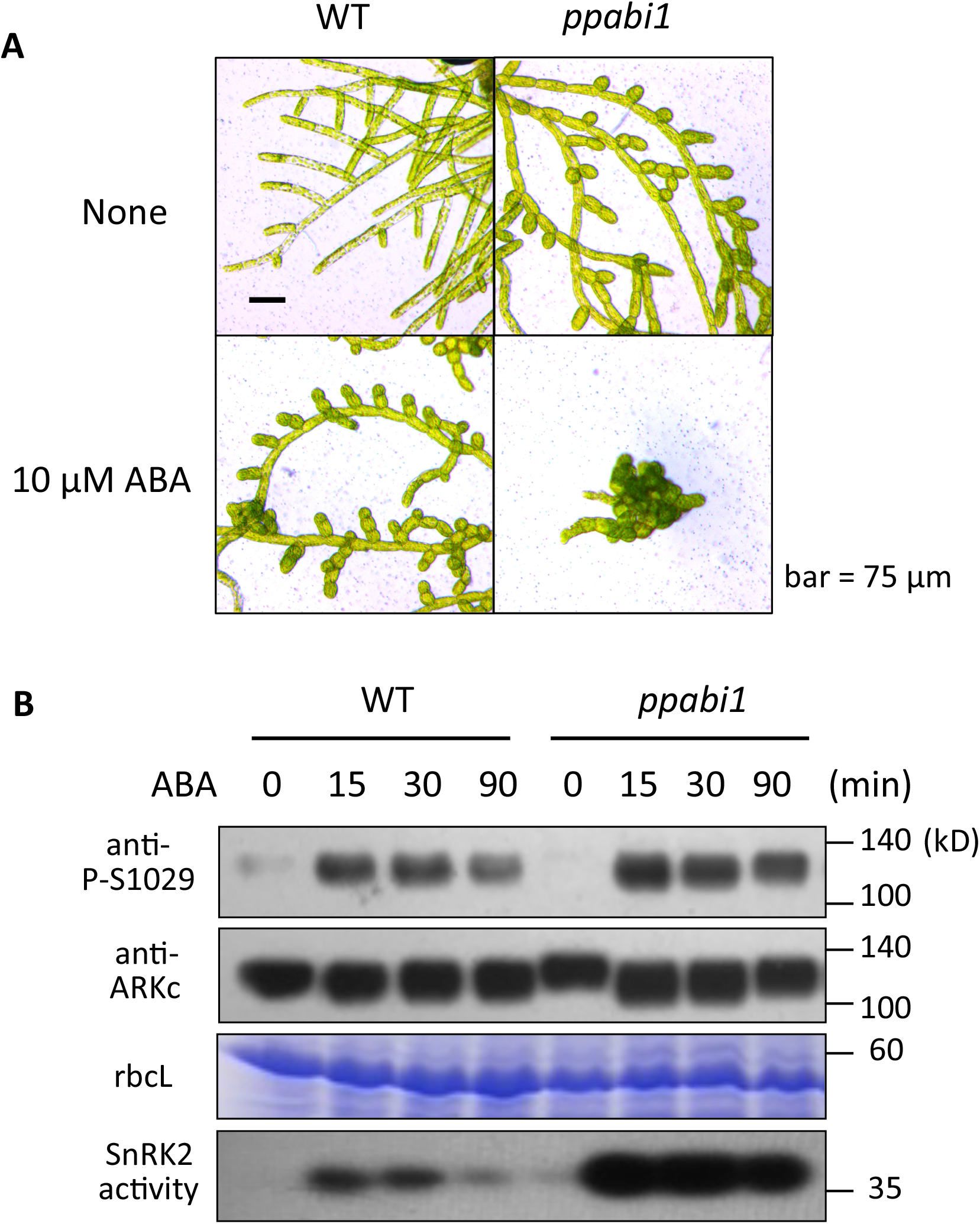
ABA response of the *ppabi1* line of *Physcomitrella patens.* (**A**) Growth responses to ABA in protonemata of WT and *ppabi1*. The protonemata were grown on a medium with or without 10 μM ABA for one week. (**B**) Effect of ABA treatment on Ser-1029 phosphorylation in WT and *ppabi1*. Protonemata were treated with 10 μM ABA for different times and used for immunoblot analysis with anti-ARKc and anti-phospho-Ser1029 (P-S1029) antibodies. Coomassie Brilliant Blue staining of the large subunit of ribulose bisphosphate calboxylase (rbcL) and SnRK2 activity analyzed by an in-gel kinase are also shown.

To determine the roles of ARK under the *ppabi1* background, we mutagenized protonemata of *ppabi1* by ultraviolet light and isolated lines that grow well on a medium containing 1 μM ABA, which severely inhibit growth of *ppabi1*. Of several isolated mutant lines, we identified ppuv4 (*ark^S1069S^*), ppuv6 (*ark^S1069L^*) and ppuv9 (*ark^S998L^*) lines (Fig. 6A, Fig. S4A) that all had missense mutations in the kinase domain. These mutant lines showed ABA-insensitive growth similar to that of AR7 (*ark^S532F^*), forming no brood cells upon ABA treatment (Fig. 6B, Fig. S4B). A comparison of ABA-induced accumulation of boiling-soluble proteins and the LEA-like 17B9 protein therein in WT, *ppabi1*, AR7 and ppuv4 indicated that AR7 and ppuv4 accumulated little of these proteins with or without ABA treatment (Fig. 6C). Furthermore, in-gel kinase assays for SnRK2 activity (Fig. 6D), freezing tolerance tests (Fig. 6E) and gene expression studies (Fig. S5) indicated that ppuv4 is insensitive to ABA. Transient reporter assays using *proEm-GUS* for determination of exogenous ABA- and ARK-dependent gene expression indicated that, while *ppabi1* shows *GUS* expression even without exogenous ABA due to its hypersensitivity, ppuv4 shows little expression with or without exogenous ABA, which can be only restored by introduction of *ARK* cDNA (Fig. 6F). These results indicated that mutations in ARK cause ABA insensitivity even under the *ppabi1* background.

**Figure 6:**
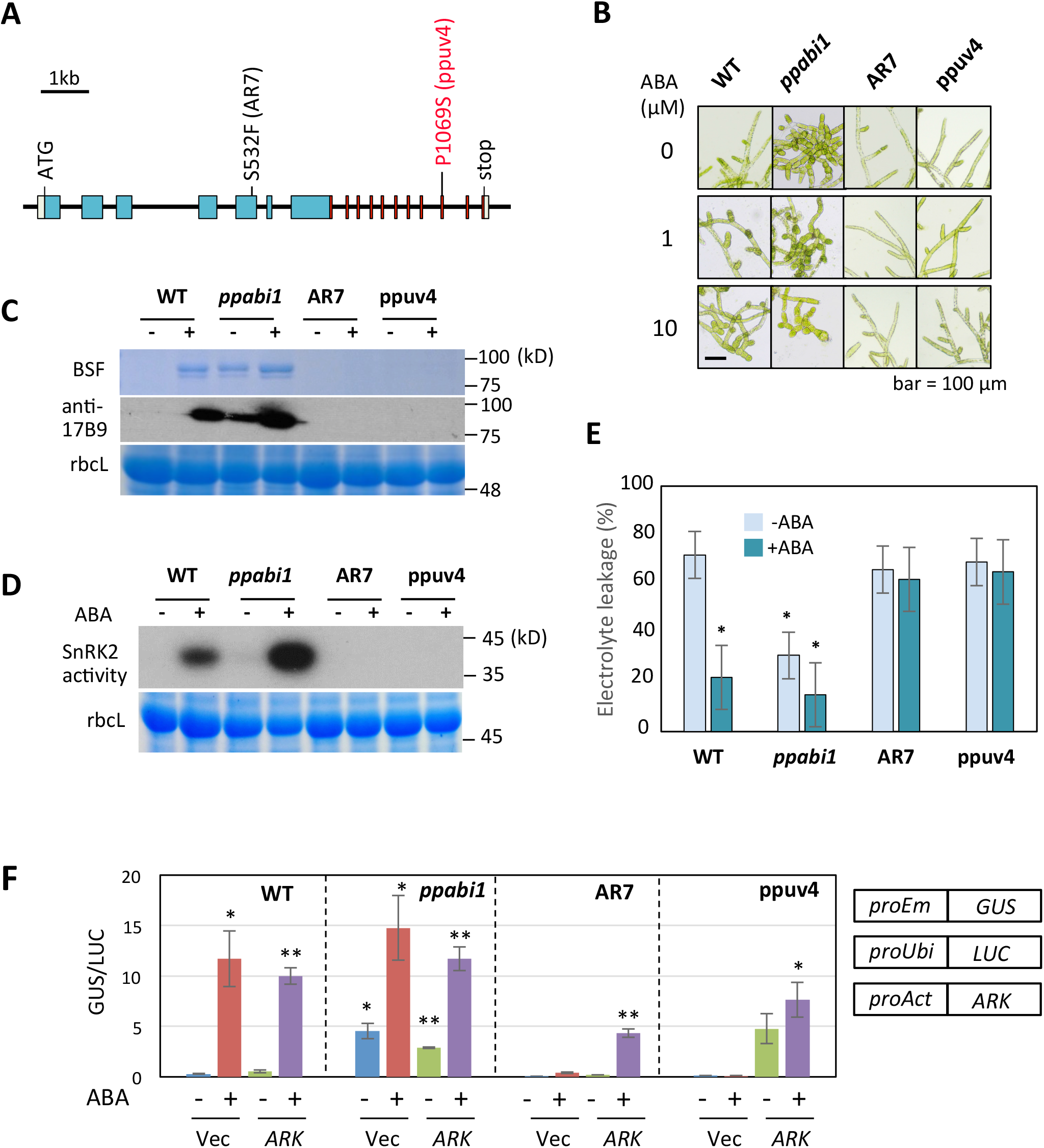
ABA response of *ppabi1 ark* lines of *Physcomitrella patens*. (**A**) Positions of mutations found in the ABA-insensitive ppuv4 line isolated by ultraviolet mutagenesis of *ppabi1*. (**B**) Effects of ABA on growth of wild type (WT), *ppabi1*, AR7 and ppuv4. (**C**) ABA-induced accumulation of boiling-soluble LEA-like proteins in WT, *ppabi1*, AR7 and ppuv4. Total soluble proteins of these lines were boiled for 1 min and centrifuged, and the boiling-soluble fraction (BSF) in the supernatant was analyzed by SDS-PAGE. The proteins were either stained with Coomassie Brilliant Blue or used for immunoblot analysis using an antibody against the LEA-like 17B9 protein. Staining of the large subunit of ribulose bisphosphate calboxylase (rbcL) of the total soluble proteins is shown as a control. (**D**) SnRK2 activity in WT and the mutant lines analyzed by in-gel kinase assays using histone IIIS as a substrate. (**E**) Freezing tolerance tests of WT and mutant lines. Protonemata were treated with or without 10 μM ABA for one day and frozen to −4°C. Electrolyte leakage (%) was measured after thawing to determine freezing injury. Error bars indicate standard error of the mean. \**p* < 0.05 in the t-test (n = 3). (**F**) Gene expression with or without 10 μM ABA treatment in WT and the mutant lines. Transient gene expression analysis was carried out using the construct of ARK fused to the rice actin promoter (*proAct*) or that without ARK (Vec). Error bars indicate standard error of the mean. \**p* < 0.05, \*\**p* < 0.01 in the t-test (n = 4) compared with the values of WT (Vec, -ABA).

## DISCUSSION

### Activation of Raf-like kinase ARK by Ser1029 phosphorylation is a key event in ABA response

We demonstrated in this study that ABA and abiotic stress responses in *P. patens* require activation of ARK, which belongs to the family of B3-Raf kinases and has been suggested to be an upstream activator of subclass III SnRK2. Previous studies indicated that nearly 90% and 80% of ABA-regulated gene expression are under the control of ARK and subgroup III SnRK2, respectively (Saruhashi *et al*., 2015; Shinozawa *et al*., 2019). Consistently, approximately 88% of ABA-increased phosphorylation was found to be ARK-dependent (Amagai *et al*., 2018), where drastic reduction in the levels of ABA-activated phosphorylation of both SnRK2 and AREB transcription factors in AR7 was observed. Our results showing correlations of Ser1029 phosphorylation with ABA-induced gene expression, stress tolerance and SnRK2 activity indicate that activation of ARK by phosphorylation is a crucial event during ABA response in *P. patens.* The role of ARK in SnRK2 activation may not be restricted to bryophytes; we previously showed that ectopic expression of three Arabidopsis B3-Raf kinases (At1G18160, At1G73660 and At4G24480) can restore ABA sensitivity of AR7 (Saruhashi *et al*., 2015). It has recently been reported that these B3-Raf kinases are necessary for activation of SnRK2 kinases during ABA and osmotic responses in Arabidopsis (Takahashi *et al*., 2020; Lin *et al*., 2020). The serine residue corresponding to Ser1029 of ARK is conserved in Arabidopsis B3-Raf kinases (Fig. S6). Considering the functional conservation of subclass III SnRK2 in *P. patens* and Arabidopsis (Chater *et al*., 2011; Shinozawa *et al*., 2019), the results of this study suggest that sequential activation of B3-Raf kinase and subclass III SnRK2 might be an evolutionarily conserved mechanism for ABA and stress responses in embryophytes. The presence of Arabidopsis B3-Raf kinases with specific functions such as CONSTITUTIVE TRIPLE RESPONSE1 (CTR1) for ethylene signaling (Kieber *et al*., 1998), ENHANCED DISEASE RESISTANCE1 (EDR1) for non-host resistance (Frye and Innes, 1998; Hiruma *et al*., 2011) and SUGAR INSENSITIVE8 (SIS8) for sugar sensing (Huang *et al*., 2014) suggests functional diversification of B3-Raf kinases in angiosperms.

### Ser1029 phosphorylation of ARK requires a functional kinase domain

The present study revealed the importance of both the kinase domain and the N-terminal region of ARK for its activation (Fig. 3E). AR144 lacking the critical Asp residue in the catalytic loop of the kinase domain showed no ABA response and Ser1029 phosphorylation, suggesting that the phosphorylation requires a functional kinase domain. This indicates that Ser1029 is not phosphorylated by other protein kinases but is phosphorylated by ARK itself (i.e., autophosphorylation). Consistently, bacterially expressed recombinant GST-ARK proteins readily undergo autophosphorylation in the activation loop (Table S3). What controls this autophosphorylation upon ABA and cold treatment *in vivo* is yet to be clarified. In animal Raf kinases, the N-terminal domains play a crucial role in regulation of the catalytic activity (Matallanas *et al*., 2011). For human c-Raf, the “autoinhibitory block” consisting of the Ras-binding domain and the C1 domain directly interacts with the C-terminal kinase domain and inhibits its catalytic activity, and upon stimulation, binding of Ras-GTP releases the kinase domain from this autoinhibition. The partially activated c-Raf undergoes autophosphorylation in the activation loop, which fully activates the kinase activity. For *P. patens* ARK, the N-terminal non-kinase region might also play a role in regulation of the kinase domain, since AR7 with S532F mutation showed little Ser1029 phosphorylation and catalytic activity toward SnRK2 (Fig. 2A and Fig. 3E). Given that the Ser1029 phosphorylation is done by autophosphorylation, activity of ARK must be controlled by the action of an unidentified negative regulatory (NR) factor that inhibits the catalytic activity of the kinase domain, possibly through interaction with a specific domain in the N-terminal region, and by ABA-dependent removal of the NR factors.

### Roles of ARK in cold responses

Many plant species can acquire tolerance to freezing upon exposure to low, non-freezing temperatures. In Arabidopsis, cold provokes expression of a number of transcripts such as those for COR/DHN with characteristics similar to LEA proteins. Expression of these transcripts is driven by CBF transcription factors, genes for which are also up-regulated by cold (Thomashow, 2010; Shi *et al*., 2018). Several distinct mechanisms for regulation of *CBF* genes have been proposed, although the molecules for cold sensing and the early signal transduction process have not been identified. It has been proposed in Arabidopsis that subclass III SnRK2 OST1 activated by cold positively regulates *CBF* gene expression by phosphorylation of the transcriptional regulators ICE1 and BTF3 (Ding *et al*., 2015; 2018). Bryophytes also undergo cold acclimation accompanied by enhancement of SnRK2 activity (Saruhashi *et al*., 2015). The fact that cold-enhanced Ser1029 phosphorylation of ARK and activation of SnRK2 were not observed in AR7 (Fig. 4B) therefore indicates the role of ARK in activation of SnRK2 for the cold response in *P. patens*. We also showed an increased accumulation of ARK polypeptide during the cold treatment (Fig. 4b), which might provide a novel mechanism for the control of ARK by cold. Interestingly, ARK transcript levels are apparently not affected by cold (Fig. 4B). It is also likely that this process does not require functional ARK, since accumulation of ARK polypeptide by cold was also observed in AR7 accumulating reduced amounts of the 17B9 protein and transcripts (Fig. 4B; Fig. S7).

### PP2C-A functions downstream to ARK

PP2C-A, which consists of nine members in Arabidopsis, is recognized as a primary regulator of SnRK2 (Umezawa *et al*., 2009; Vlad *et al*., 2009). The role of PP2C-A in negative regulation of ABA signaling is likely to be conserved in both mosses and liverworts consisting of the basal lineage of embryophytes, because overexpression of PP2C-A genes of these plants caused abolishment of ABA-responsive gene expression and loss of abiotic stress tolerance (Komatsu *et al*., 2009; Tougane *et al*., 2010). Furthermore, the fact that the *ppabi1* plants still responded to ABA and could activate SnRK2 (Komatsu *et al*., 2013) indicates the presence of PP2C-A-independent ABA-response mechanisms in bryophytes. Consistently, results of phosphoproteome analysis of *P. patens* indicated that only a small portion (23 out of 143) of ABA-stimulated phosphoproteins detected in WT were hyperphosphorylated in *ppabi1*, while 126 of those were dependent on ARK (Amagai *et al*., 2018).

The present study on ppuv4 (*ppabi1 ark*) demonstrated that, with or without PP2C-A, *P. patens* cannot respond to ABA without functional ARK (Fig. 6A-F). Abolishment of SnRK2 activity in ppuv4 as in AR7 indicates that SnRK2 cannot undergo regulation by PP2C-A unless it has been pre-activated by ARK (Fig. 6D). Furthermore, our results suggested that PP2C-A may not be a primary regulator of ARK phosphorylation (Fig. 5B). Figure 7 and Figure S8 illustrate working models for the control of SnRK2 in bryophytes, focusing on the role of ARK for positive regulation and PP2C-A for subsequent negative regulation. These models are supported not only by abolishment of ABA responses in ppuv4 but also by the results of our experiment with *MpPYL1* (Fig. S9). MpPYL1 is a major ABA receptor in vegetative tissues of *M. polymorpha* and has recently been shown to have ABA-independent inhibitory activity towards PP2C-A (Jahan *et al*., 2019; Sun *et al*., 2019). Our experiment indicated that transient overexpression of *MpPYL1* enhances the activity of the *GUS* reporter gene fused to the ABA-inducible *Em* promoter without ABA treatment, but such enhancement was not observed in AR7 with or without ABA (Fig. S9), suggesting that inhibition of PP2C-A by PYR/PYL/RCAR causes activation of the promoter only in the presence of functional ARK. Our previous analysis indicated spatial separation of ARK and PP2C-A in bryophytes; The localization of the GFP-fusion protein of ARK was mainly in the cytoplasm and partly in the endoplasmic reticulum, while that of PP2C-A was only in the nucleus (Komatsu *et al*., 2009; Tougane *et al*., 2010; Saruhashi *et al*., 2015). The working models we proposed for bryophytes in this study represent the basic ABA response machinery prototypal to that in angiosperms possessing diverse PP2C-A molecules. Evolutionary diversification of PP2C-A molecules in angiosperms resulted in localization of some PP2C-A molecules in the nucleus for regulation of transcription factors, while others are in the cytoplasm for regulation of angiosperms-specific substrates such as plasma membrane-localized ion channels. Takahashi *et al*. (2020) hypothesized from the results of *in vitro* and *Xenopus* oocyte reconstitution assays that Arabidopsis B3-Raf kinases play a role in reactivation of SnRK2 that had been dephosphorylated by PP2C-A for regulation of the AKS1 transcription factor and the S-type ion channel SLAC1. We do not rule out the possibility of a role of *P. patens* ARK in reactivation of SnRK2 because *P. patens* SnRK2 is localized in both the cytoplasm and nucleus (Saruhashi *et al*., 2015), but verification of its possible role requires further investigation.

**Figure 7:**
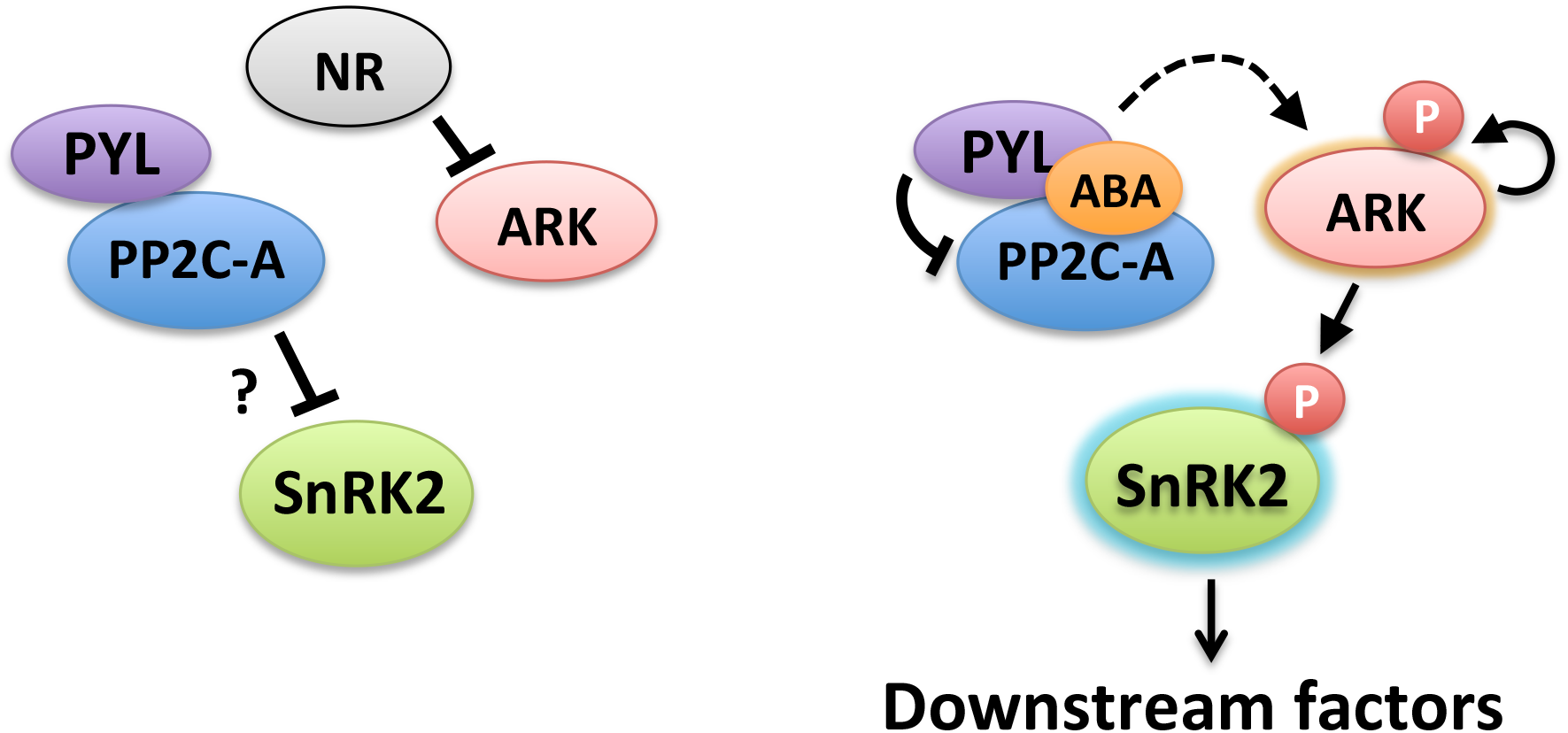
Working models showing ARK-mediated ABA responses in *Physcomitrella patens*. ABA stimulates ARK autophosphorylation and inhibits PP2C-A, causing activation of SnRK2 regulating downstream factors, in which phosphorylation of SnRK2 by ARK proceeds negative regulation by PP2C-A. ABA-stimulated activation of both ARK and SnRK2 occurs without PP2C-A, suggesting the presence an unidentified negative regulator (NR). Details of these models are shown in Fig. S8.

## MATERIALS AND METHODS

### Chemicals and plant materials

Chemicals were purchased from Wako Jun-yaku (Osaka, Japan) unless otherwise stated. Abscisic acid was from Sigma (A4906, St. Louis, MO, USA). Culture and ABA, osmotic and cold treatments of protonemata of *P. patens* wild type (WT) and mutant lines were carried out as described previously (Minami *et al*., 2003; 2005).

### Transient reporter assays

*P. patens* protonemata cultured on cellophane-overlaid BCDAT agar medium for 5 days were used for particle bombardment (Marella *et al.*, 2006). The protonema cells were bombarded with DNAs of the *proEm*-*GUS* reporter construct (Marcotte *et al*., 1989) and the *proUbi-LUC* reference construct (Bruce *et al*., 1989) with or without *ARK* effector constructs by using the PDS-1000He particle delivery system (Bio-Rad, Hercules, CA). The bombarded cells were incubated in a medium with or without ABA and were used for GUS and LUC assays (Bruce *et al*., 1989; Jefferson *et al*., 1989; Marella *et al.*, 2006; Komatsu *et al*., 2009).

### Generation of transgenic *P. patens*

*P. patens* protonemata were subjected to PEG-mediated transformation according to the protocol described by Nishiyama *et al*. (2000). Constructs of ARK-green fluorescent protein (ARK-GFP) without mutation and that with mutations in Ser532 to Phe (ARK^S532F^-GFP), Ser1029 to Ala (ARK^S1029A^-GFP), Ser1030 to Ala (ARK^S1030A^-GFP) and Ser1029/Ser1030 to Ala/Ala (ARK^S1029A/S1030A^-GFP) driven by the rice actin promoter were made using the pFL-Not vector (Saruhashi *et al*., 1995).

### Mutant screening

Partially homogenized protonemata spread on a cellophane-overlaid agar medium were cultured for two days prior to irradiation of ultraviolet light under conditions described by Minami *et al*. (2006). After incubation in the dark at 25°C for one day, the protonemata were cultured under continuous light for 1 or 2 days at 25°C. They were then transferred with cellophane onto a medium containing 10 μM ABA for WT and onto a medium containing 1 μM ABA for *ppabi1*.

### Protein gel electrophoresis and immunoblot analysis

For protein extraction, protonemata were homogenized in a buffer containing 50 mM Tris-Cl (pH 8.0), 150 mM NaCl, 1% Triton X-100, 25 mM NaF, 2 mM dithiothreitol, 1 mM o-vanadate, and 50 mM β-glycerophosphate and a 1/100 volume of proteinase inhibitor mixture (P9599; Sigma, MO) on ice. After centrifugation at 14,000 × g for 10 min at 4 °C, the supernatants were used as protein samples for electrophoresis. The proteins (30 μg) were electrophoresed on 8% SDS-polyacrylamide gel and blotted onto a polyvinylidene fluoride membrane for immunoblot analysis. After blocking with 1% bovine serum albumin (for the anti-phosphopeptide antibody) or 3% skim milk (for other antibodies) in TBS-T (25 mM Tris-Cl, pH 7.5, 150 mM NaCl and 0.05%(w/v) Tween-20), the proteins were reacted with the following primary antibodies: anti-GFP (MBL, Nagoya, Japan), anti-ARK C-terminal 15 amino acids (LGGTPKSGLSDRDL) (anti-ARKc) (Saruhashi *et al*., 2015) and anti-Ser1029-phosphorylated peptide (FLTSKpSSAGTPEWMAPE) (anti-P-Ser1029). The anti-P-Ser1029 antibody raised in rabbits was affinity-purified using the same phosphopeptide and then cross-absorbed against the dephosphorylated GST-ARK protein to remove antibody that might recognize non-phosphorylated ARK. The membrane that had been reacted with a primary antibody was reacted with a 1:10000 dilution of horseradish peroxidase-conjugated secondary antibody (MBL, Nagoya, Japan). The membrane was then immersed in chemiluminescence reagent (Chemi-Lumi One, Nacalai Tesque, Kyoto, Japan) and exposed to X-ray film for signal detection.

### Analysis of phosphopeptides

Protein extracts were prepared from protonemata of AR7 expressing ARK-GFP constructs using the buffer used for immunoblot analysis. Immunoprecipitation was carried out using the μMACS GFP Tagged Protein Isolation Kit (Miltenyi Biotech). Protein samples were electrophoresed using 8% SDS polyacrylamide gel and stained with Coomassie Brilliant Blue, and stained bands corresponding to ARK-GFP near 150 kilodaltons were excised. After proteolytic enzyme digestion, the phosphopeptides were analyzed by nano-flow reverse-phase liquid chromatography followed by tandem MS using a Q Exactive Hybrid Quadrupole-Orbitrap Mass Spectrometer (Thermo Fisher Scientific).

### Extraction of total RNA, reverse transcription (RT)-PCR and RT-PCR analysis

RNA extraction, first-strand cDNA synthesis and PCR reaction were carried out as described previously (Saruhashi *et al*., 2015). The RNA was reverse-transcribed with oligo-dT and used for semi-quantitative PCR and SYBR Green-based quantitative PCR. Sequences of oligonucleotide primers are listed in Table S1.

### In-gel kinase assay for SnRK2 activity

An in-gel kinase assay was carried out according to the protocol described by Suzuki and Shinshi (1995). Protein extracts used for immunoblot analysis were subjected to electrophoresis using a 10% SDS-polyacrylamide gel containing 0.5 mg/mL histone IIIS (Sigma-Aldrich) as a substrate. The gel was successively washed for 30 min twice in buffer I (50 mM Tris-Cl, pH 8.0 and 20% isopropanol) and buffer II (50 mM Tris-Cl, pH 8.0 and 5 mM 2-mercaptoethanol), denaturation buffer (buffer II containing 6 M guanidine hydrochloride), and renaturation buffer (buffer II containing 0.04% Tween 40) at room temperature. The gel was then incubated in renaturation buffer at 4°C for 16 h and again washed once in renaturation buffer at room temperature for 30 min. After incubation in buffer III (40 mM HEPES-KOH, pH 7.5, 15 mM MgCl_2_, 0.1 mM ethylene glycol tetraacetic acid and 2 mM dithiothreitol) for 30 min at room temperature, the gel was reacted with 50 μM [γ-^32^P] ATP (1.5 TBq/mmol) in buffer III for 1 h at 25°C. After removal of the reaction solution, the gel was washed for 30 min for at least six times with washing solution (5% trichloroacetic acid and 1% sodium pyrophosphate) before drying and exposure to X-ray film.

### Tests for freezing tolerance

Protonema tissues placed in a glass test tube containing 0.5 mL of distilled water were set in a programmable cooling bath. The tissues were seeded with ice at −1°C, kept at −1°C for 50 min, and then cooled at a rate of −2.4°C h^−1^ to desired temperatures. Electrolyte leakage from the damaged tissues was determined after thawing by measurement of conductivity in the water and represented as the percentage against conductivity of total ions released by subsequent boiling.

## ACKNOWLEDGEMENTS

We thank Yasuo Niwa for GFP(S65T) (Niwa, 2003), Ralph Quatrano for *proEm-GUS*, and Tuan-hua David Ho for *Ubi-LUC* constructs. We also thank Masashi Saruhashi and Marina Fujita for technical assistance.

## SUPPORTING MATERIAL

**Table S1:** Oligonucleotide primers used for gene expression studies.

**Table S2:** Analysis of phosphopeptides immunoprecipitated from AR7 expressing ARK-GFP with or without mutations in Ser1029 and Ser1030.

**Table S3:** Autophosphorylation in the peptide SSAGTPEWMAPEVLR in the activation loop of recombinant GST-ARK.

**Figure S1:** Growth responses to ABA of transgenic AR7 lines expressing various ARK-GFP constructs.

**Figure S2:** Detection of Ser1029 phosphorylation of native ARK.

**Figure S3:** Effects of hyperosmotic mannitol on ARK Ser1029 phosphorylation.

**Figure S4:** Growth responses to ABA of three ppuv mutants isolated by ultraviolet light mutagenesis of *ppabi1*.

**Figure S5:** ABA-induced gene expression of *LEA*-like transcripts in wild type (WT), *ppabi1*, AR7 and ppuv4.

**Figure S6:** Amino acid sequence alignment in the activation loops of ARK and Arabidopsis B3-Raf kinases.

**Figure S7:** Expression of *ARK* and *LEA*-like *17B9* transcripts during cold acclimation in WT and AR7.

**Figure S8:** Details of the working model shown in Figure 7.

**Figure S9:** Effect of MpPYL1 on ABA-inducible promoter in WT and AR7.

